# Reinforcement of Fibrillar Collagen Hydrogels with Bioorthogonal Covalent Crosslinks

**DOI:** 10.1101/2025.04.13.648560

**Authors:** Lucia G. Brunel, Chris M. Long, Fotis Christakopoulos, Betty Cai, Narelli de Paiva Narciso, Patrik K. Johansson, Diya Singhal, Annika Enejder, David Myung, Sarah C. Heilshorn

## Abstract

Bioorthogonal covalent crosslinking stabilizes collagen type I hydrogels, improving their structural integrity for tissue engineering applications with encapsulated living cells. The chemical modification required for crosslinking, however, interferes with the fibrillar nature of the collagen, leading instead to an amorphous network without fibers. We demonstrate an approach to perform bioconjugation chemistry on collagen with controlled localization such that the modified collagen retains its ability to self-assemble into a fibrillar network, while also displaying functional groups for covalent crosslinking with bioorthogonal click chemistry. The collagen matrix is formed through a sequential crosslinking process, in which the modified collagen first physically assembles into fibers and then is covalently crosslinked. This approach preserves the fibrous architecture of the collagen, guiding the behavior of encapsulated human corneal mesenchymal stromal cells, while also reinforcing fibers through covalent crosslinks, strengthening the stability of the cell-laden collagen hydrogel against cell-induced contraction and enzymatic degradation.

## 1. Introduction

As the most abundant protein in the human body, collagen is a commonly used biomaterial for tissue engineering and regenerative medicine applications. In particular, 90% of connective tissue is composed of collagen type I, which can be easily extracted from tissues and purified to minimize contamination from other extracellular matrix (ECM) components.^1,2^ Collagen type I is a triple helical molecule that physically self-assembles into fibrils in an aqueous medium with neutral pH, forming a hydrogel.^3^ Collagen products such as dermal fillers have already been approved for medical and cosmetic uses in humans by the U.S. Food and Drug Administration.^4^

In addition to the applications of acellular collagen hydrogels, collagen hydrogels have also been explored as suitable 3D scaffolds for the encapsulation of living cells. Collagen hydrogels guide essential cell behaviors by presenting both biochemical cues like ligand-binding sites and biophysical cues like fibrous hierarchical architectures.^5–9^ While the collagen microenvironment can support cell viability, spreading, migration, and differentiation, collagen hydrogels commonly suffer from poor mechanical stability.^2^ Forces exerted on collagen networks by encapsulated, contractile cells such as fibroblasts or mesenchymal stromal cells (MSCs) can cause severe, undesired shrinkage and compaction of the tissue-engineered constructs.^10–15^ Furthermore, collagen hydrogels are susceptible to enzymatic degradation, particularly by collagenase, a protease commonly secreted by cells that plays a role in many pathological conditions including wound healing.^14–16^ If the enzymatic degradation of the collagen hydrogel occurs at a faster rate than the production of nascent ECM from encapsulated cells, the engineered tissue can weaken and fall apart.^17^ Therefore, the use of collagen hydrogels as a basis for *in vitro* tissue models or *in vivo* transplantation requires suitable stability of the tissue-engineered construct against contractile forces and proteases produced by encapsulated cells.

Conventional approaches to tune the stability of collagen hydrogels have historically included physical processes such as dehydrothermal treatment (DHT) or chemical processes such as crosslinking with glutaraldehyde, 1-ethyl-3-(3-dimethyl aminopropyl) carbodiimide hydrochloride (EDC), or N-hydroxysuccinimide (NHS).^18,19^ While these techniques can efficiently strengthen and stabilize acellular collagen hydrogels, their cytotoxicity prohibits the inclusion of cells within the collagen matrix during the crosslinking process,^20,21^ instead requiring cells to be seeded onto the top of the scaffold after crosslinking.^22–24^ Cytocompatible crosslinking strategies have therefore been developed to enable the direct inclusion of cells within the collagen, allowing for homogenous distribution of cells.^25–27^ In particular, strain-promoted azide-alkyne cycloaddition (SPAAC) is a copper-free click chemistry that proceeds at physiological temperature and pH.^28^ The bioorthogonal nature of SPAAC allows the chemical reaction to selectively occur in biological environments without interfering with living cells or other biomolecules.^27^ To crosslink collagen with SPAAC chemistry, a solution of collagen is modified with a SPAAC reactive group (*i.e.*, either azides or strained alkynes), mixed with the desired cell type, and then crosslinked with the complementary SPAAC reactive group, often displayed on a hydrophilic molecule such as polyethylene glycol (PEG).^13–15^

While SPAAC crosslinking successfully improves the structural stability of collagen hydrogels, the chemical modification process has previously disrupted the ability of the collagen molecules to self-assemble into fibrils.^13–15,29,30^ Therefore, previous SPAAC-crosslinked collagen networks have all been amorphous, without the presence of fibers.^13–15,29,30^ Since the native, fibrous architecture of collagen strongly affects cellular interactions, a recent approach explored the design of an interpenetrating network (IPN) combining an amorphous SPAAC-crosslinked collagen network with a fibrous, physically-assembled (PHYS) collagen network.^14^ In this current work, we hypothesized that we could instead alter the physical state of the collagen during the bioconjugation process to selectively modify the collagen fibril surface with azides, such that the collagen not only is amenable to SPAAC crosslinking but also retains its ability to self-assemble into fibrils. We demonstrate the microstructural differences of collagen modified with azides in either the solution-phase, leading to an amorphous SPAAC (a-SPAAC) collagen network, or the gel-phase, leading to a fibrillar SPAAC (fib-SPAAC) collagen network. By leveraging fib-SPAAC collagen, viability and spreading of encapsulated human cells are facilitated by the collagen fibers, while increased stability of the bulk hydrogel is facilitated by the bioorthogonal, covalent SPAAC crosslinking. These results indicate a chemical strategy to enhance cell-laden collagen hydrogels with both biophysical functionality and structural integrity.

## 2. Materials and Methods

### 2.1. Synthesis of collagen-azide

#### Collagen-azide bioconjugation in solution-phase

Bovine type I atelocollagen solution (10 mg/mL, Advanced BioMatrix) was first brought to a concentration of 8 mg/mL collagen and neutral pH of ∼7.4 according to the manufacturer’s protocol using 10X phosphate buffered saline (PBS, Millipore), 1.0 M sodium hydroxide (NaOH, Sigma), and ultrapure deionized water (Millipore). To keep the collagen in solution, all neutralization steps and the bioconjugation reaction were conducted on ice or at 4 °C. The primary amines of collagen were modified with azide groups using NHS ester chemistry. A 100 mg/mL solution of azido-PEG_4_-NHS ester (BroadPharm) in dimethyl sulfoxide (DMSO, Fisher) was added to the neutralized collagen solution at 2 molar equivalents relative to primary amines in collagen. The solution was rotated at 4 °C for 2 h, dialyzed overnight at 4 °C with a Slide-A-Lyzer dialysis kit (3.5 kDa-MWCO, Thermo Scientific) against pre-cooled 1X PBS, and stored at 4 °C before use.

#### Collagen-azide bioconjugation in gel-phase

Bovine type I atelocollagen solution (10 mg/mL, Advanced BioMatrix) was first brought to a concentration of 8 mg/mL collagen and neutral pH of ∼7.4 according to the manufacturer’s protocol using 10X PBS (Millipore), 1.0 M NaOH (Sigma), and ultrapure deionized water (Millipore). 5 mL of collagen solution were transferred into the bottom of a 10-cm diameter Petri dish (Falcon) and incubated at 37 °C for 1 h to form a hydrogel. The primary amines of collagen were modified with azide groups using NHS ester chemistry, by adapting a protocol for modifying collagen hydrogels.^31^ 10 mL of 50 mM borate buffer (pH = 9.0) were first added atop the collagen hydrogel and incubated at room temperature (RT) for 15 min. The borate buffer was then replaced with a solution of 5 mL of the borate buffer containing azido-PEG_4_-NHS ester (BroadPharm; dissolved in DMSO) at 2 molar equivalents relative to primary amines in collagen and incubated at RT for 1 h while rocking. After 1 h, the borate buffer with azido-PEG_4_-NHS ester was aspirated. 10 mL of 50 mM tris buffer (pH = 7.5) were added and incubated at RT for 10 min to quench the reaction. The hydrogel was then washed with PBS (6 washes, 30 min each) at RT while rocking. After the final PBS wash was removed, the collagen was resolubilized by adding 500 µL of 200 mM hydrochloric acid (HCl, Sigma-Aldrich) and incubating at 4 °C for 1 h while rocking. The solution was then transferred using a positive-displacement pipet into a glass scintillation and stirred overnight at 4 °C. After the hydrogel had fully returned to solution, it was dialyzed overnight at 4 °C with a Slide-A-Lyzer dialysis kit (3.5 kDa-MWCO, Thermo Scientific) against pre-cooled 20 mM acetic acid (Fisher) and then stored at 4 °C before use.

#### Degree of functionalization characterization

The degree of functionalization for collagen-azide modified in either the solution-phase or gel-phase was determined using a 2,4,6-Trinitrobenzene Sulfonic Acid (TNBSA) assay (Thermo Scientific) to quantify free amino groups, following the instructions from the manufacturer. Unmodified collagen was used as the control.

### 2.2. Preparation of hydrogels

#### Physically self-assembled (PHYS) collagen hydrogel

Bovine type I atelocollagen solution (10 mg/mL, Advanced BioMatrix) was first brought to a concentration of 8 mg/mL collagen and neutral pH of ∼7.4 according to the manufacturer’s protocol using 10X PBS (Millipore), 1.0 M NaOH (Sigma), and ultrapure deionized water (Millipore). The neutralized solution was diluted to 6 mg/mL collagen with cold 1X PBS. To form the PHYS collagen hydrogel, the neutralized collagen solution was transferred to 8-well chambered coverslips (Ibidi) and then incubated at 37 °C for 1 h for gelation.

#### PHYS + amorphous Strain-Promoted Azide-Alkyne Cycloaddition (a-SPAAC) collagen hydrogel

A solution with a concentration of 4 mg/mL unmodified PHYS collagen and 2 mg/mL collagen-azide (sol-phase modified) was prepared on ice and mixed thoroughly. The solution was transferred to 8-well chambered coverslips (Ibidi) and incubated at 37 °C for 1 h for gelation through collagen physical self-assembly. A polyethylene glycol-dibenzocyclooctyne (PEG-DBCO, 4-arm, 10 kDa, Creative PEGworks) crosslinker dissolved in 1X PBS was then added atop each hydrogel for an effective concentration of 4 mg/mL PEG-DBCO. Finally, the PHYS + a-SPAAC hydrogel was incubated again at 37 °C for 1 h for the SPAAC reaction to proceed, forming the interpenetrating network.

#### PHYS + fibrillar Strain-Promoted Azide-Alkyne Cycloaddition (fib-SPAAC) collagen hydrogel

For simultaneous gelation, the two gelation mechanisms (*i.e.*, physical self-assembly and SPAAC crosslinking) were allowed to proceed simultaneously. A solution of 4 mg/mL unmodified PHYS collagen, 2 mg/mL collagen-azide (gel-phase modified, resolubilized), and 4 mg/mL PEG-DBCO was mixed well on ice. The solution was then transferred to 8-well chambered coverslips (Ibidi) and incubated at 37 °C for 1 h. For sequential gelation, the physical self-assembly was allowed to proceed before the SPAAC crosslinking reaction. First, a solution of 4 mg/mL unmodified PHYS collagen and 2 mg/mL collagen-azide (gel-phase modified) was transferred to 8-well chambered coverslips (Ibidi) and incubated at 37 °C for 1 h for gelation through physical self-assembly. Then, the PEG-DBCO crosslinker dissolved in PBS was added atop each hydrogel for an effective concentration of 4 mg/mL PEG-DBCO and incubated again at 37 °C for the SPAAC reaction to proceed.

### 2.3. Circular dichroism (CD) spectroscopy

Collagen samples were diluted to 0.2 mg/mL in 1X PBS. Neutralized, unmodified collagen was used as a positive control. Neutralized, heat-denatured collagen that had been incubated at 90 °C for 2 h was used as a negative control. CD spectroscopy was conducted using a Jasco J-815 spectropolarimeter. Samples were loaded into a cuvette (Hellma) with a 1-mm pathlength, and the CD spectra were acquired at 20 °C within a wavelength interval of 190-250 nm. The scan speed was set at 50 nm/min with a pitch of 1 nm. Each spectrum was averaged across 3 scans.

### 2.4. Rheometry

Rheological characterization was performed using an ARG2 stress-controlled rheometer (TA Instruments). All measurements were confirmed to be within the linear viscoelastic regime of the materials. Tests comparing the physical self-assembly of the collagen-azide materials were conducted with a cone-and-plate geometry (20-mm diameter, 1° cone). The solutions, initially kept on ice, were pipetted onto the rheometer. The gelation kinetics were then measured with a temperature sweep between 4-37 °C with a 3 °C/min heating rate followed by a time sweep with 1 rad/s angular frequency and 1% strain. Tests comparing the shear moduli and stress relaxation time of PHYS + fib-SPAAC with and without the SPAAC crosslinker (*i.e.*, PEG-DBCO) were conducted with a parallel plate geometry (8-mm diameter). First, a solution of 4 mg/mL unmodified PHYS collagen and 2 mg/mL collagen-azide (gel-phase modified) was gelled within silicone molds (8-mm diameter). For the condition without the SPAAC crosslinker, the hydrogels were ready for measurement. For the condition with the SPAAC crosslinker, the hydrogels were first exposed to PEG-DBCO at an effective concentration of 4 mg/mL PEG-DBCO for 1 h at 37 °C before the measurement. Hydrogels were removed from their molds and placed onto the rheometer stage. Frequency sweeps were conducted between 0.1-100 rad/s with 1 % strain at 37 °C. Stress relaxation measurements were conducted with 10 % strain at 37 °C.

### 2.5. Enzymatic degradation assay

Accelerated enzymatic degradation assays were conducted on PHYS + fib-SPAAC collagen with varying concentrations of PEG-DBCO crosslinker, ranging from 0-32 mg/mL. The hydrogels were prepared in Eppendorf tubes and exposed to 5 mg/mL of collagenase (Gibco) dissolved in PBS. The samples were incubated at RT on an orbital shaker. At each timepoint, the collagenase solution was removed, the wet weight of the sample was recorded, and fresh collagenase was added.

### 2.6. Cell culture and encapsulation

Human corneal mesenchymal stromal cells (MSCs) were isolated from a donor cornea (Lions Eye Institute for Transplant and Research) using established protocols.^32^ Cell growth medium was prepared with the following formulation: 500 mL MEM-Alpha (Corning), 50 mL fetal bovine serum (Gibco), 5 mL GlutaMax (Gibco), 5 mL non-essential amino acids (Gibco), and 5 mL antibiotic-antimycotic (Gibco). The growth medium was changed every other day. The cells were passaged upon reaching 80% confluency using 0.05% Trypsin-EDTA (Gibco) for trypsinization. Cells were used for experiments between passages 7-9. For all *in vitro* cell experiments, cells were encapsulated within PHYS or PHYS + fib-SPAAC collagen at a density of 5 × 10^!^ cells/mL. The cell-laden materials were cast into silicone molds (4-mm diameter) for the contraction assay and into 18-well glass bottom chamber slides (Ibidi) for the cell viability and spreading assays.

### 2.7. Second harmonic generation (SHG) microscopy and image analysis of collagen fibrils

Fibrillar collagen (distinct from amorphous collagen) was visualized using second harmonic generation (SHG) microscopy on an inverted Nikon Ti2-E microscope with a C2 confocal scanner. A Nikon CFI Apochromat TIRF 100X oil-immersion objective was used for imaging. A sliding mirror (Optique Peter) allowed for easy alternation between confocal fluorescence and SHG excitation modalities. For SHG, a picosecond-pulsed laser system (picoEmerald S, APE America Inc.) emitting 2 ps pulses at an 80 MHz repetition rate and with a 10 cm^-1^ spectral bandwidth was used. The system consists of a tunable optical parametric oscillator (700-960 nm) pumped at 1031 nm by an ytterbium fiber laser. The oscillator was set to 797 nm, generating an SHG signal at 398.5 nm. The emitted signal was routed through optical filters (BrightLine 400/12, Thorlabs 390/18, and Thorlabs FESH0500) before detection by a photomultiplier tube (Hamamatsu R6357). The laser power at the sample was 35 mW.

To facilitate imaging, hydrogel samples were removed from their molds and placed into 2-well chamber slides (Ibidi, 1.5H glass coverslip) with ∼500 μL PBS to avoid drying. Nine distinct regions were imaged per sample, each captured as a 21-slice z-stack (1 μm slice spacing, total depth 20 μm). To minimize boundary effects, the midpoint of each z-stack was positioned at least 30 μm deep into the gel surface. Images were acquired at 1024 × 1024 pixels (75.3 × 75.3 nm/pixel) with a dwell time of 10.8 μs/pixel.

The SHG z-stacks were processed in FIJI,^33^ where an edge-detection filter and a 3D Gaussian blur (σ = 1) were first applied, followed by Otsu’s thresholding method^34^ to generate a binary mask that isolated collagen fibrils. Fibril thicknesses were then measured using the “Local Thickness” plugin. Fibril contour lengths, persistence lengths, mesh sizes, and fibril counts were determined using a 3D fiber-tracing algorithm in MATLAB, adapted from Rossen *et al*.^35^ A 3D Gaussian PSF was rescaled to match our voxel size, and the following parameters were selected to identify fibrils: length-prioritization of [30 15 10], cone angle of 30°, and stop factor of 0.5.

### 2.8. Fluorescence microscopy of SPAAC collagen network

To visualize the SPAAC collagen network (fibrillar as well as non-fibrillar), the PHYS + a-SPAAC collagen and PHYS + fib-SPAAC collagen hydrogels were first prepared as described in Section 2.2. The hydrogels were then incubated in 30 µM Alexa Fluor 488-DBCO (Click Chemistry Tools) in PBS with 1 wt% bovine serum albumin (Sigma-Aldrich) for 1 h at 37 °C. The hydrogels were washed 4 times with PBS to remove unreacted dye. Fluorescence imaging was conducted on the same microscope platform as that used for SHG microscopy (Section 2.7): Nikon, Ti2-E equipped with a C2 confocal scanning head and a Nikon CFI Apochromat TIRF 100XC oil immersion objective. This enabled simultaneous fluorescence and SHG measurements and, hence, the distinction of fibrous versus amorphous collagen architectures.

### 2.9. Microscopy and image analysis of cells within gels

The assessment of corneal MSC viability within hydrogels was conducted with a Live/Dead assay using calcein AM and ethidium homodimer-1 (Life Technologies), following instructions from the manufacturer. The cells were imaged with a STELLARIS 5 confocal microscope (Leica) with a 10X air objective, and the number of live and dead cells were quantified using FIJI. Cell viability was calculated as the number of live cells divided by the total number of cells. The aspect ratio (*i.e.*, ratio of the major to minor axis length) of cells was determined for living cells (*i.e.*, calcein AM-positive cells) using CellProfiler with two-class Otsu thresholding, hole removal, watershed algorithm to separate touching cells, and removal of objects with surface areas <50 µm^2^. The major axis length of the cell is the length of the major axis of the ellipse that has the same normalized second central moments as the region. The minor axis length of the cell is the length of the minor axis of the ellipse that has the same normalized second central moments as the region. For both cell viability and cell aspect ratio, analyses were conducted on 3 z-stack images (110 µm height) taken in different regions of each replicate sample.

The contraction of corneal MSC-laden hydrogels was tracked over 5 days in culture using images taken with an epifluorescent microscope (Leica Microsystems, THUNDER Imager) with a 2.5X air objective in bright field mode. Hydrogels were imaged within the original 4-mm diameter molds in which they were fabricated to show the changes in size. The areas of each sample were quantified using FIJI.

Images of the interactions between calcein AM-stained corneal MSCs and the fibrillar matrix of PHYS + fib-SPAAC collagen hydrogels were taken with a STELLARIS 5 confocal microscope (Leica) by combining fluorescence and confocal reflectance imaging modalities. Images were taken with a 40X oil immersion objective.

### 2.10. Statistical analysis

Statistical significance testing for all data was performed using the GraphPad Prism (Version 10) software package. Details of the sample sizes and statistical tests conducted for each plot are provided in the figure captions. Significance cutoffs were defined as follows: not significant (ns, *p* > 0.05), * (*p* < 0.05), ** (*p* < 0.01), *** (*p* < 0.001), and **** (*p* < 0.0001).

## 3. Results and Discussion

### 3.1. Influence of collagen chemical modification on hydrogel microstructure

For collagen to be covalently crosslinked with SPAAC chemistry, the collagen must first be chemically modified with either azides or strained alkynes.^13–15,29,30,36^ We chose to functionalize collagen with azides due to the higher hydrophilicity of azides compared to strained alkynes.^37^ This choice alleviates concerns of self-aggregation from pi-stacking or hydrophobic interactions in collagen modified with strained alkynes. To synthesize the collagen-azide (**Figure 1**), we applied NHS ester reaction chemistry to bovine type I atelocollagen. The NHS ester-activated azide molecules react with the protein’s primary amines,^38^ forming an amide bond that conjugates the azides onto the collagen. Since atelocollagen does not contain the N- and C-terminal telopeptides, which are commonly considered responsible for the immunogenicity of collagen,^39,40^ the primary amines of atelocollagen are primarily present in the sidechains of lysine amino acid residues.

**Figure 1.**
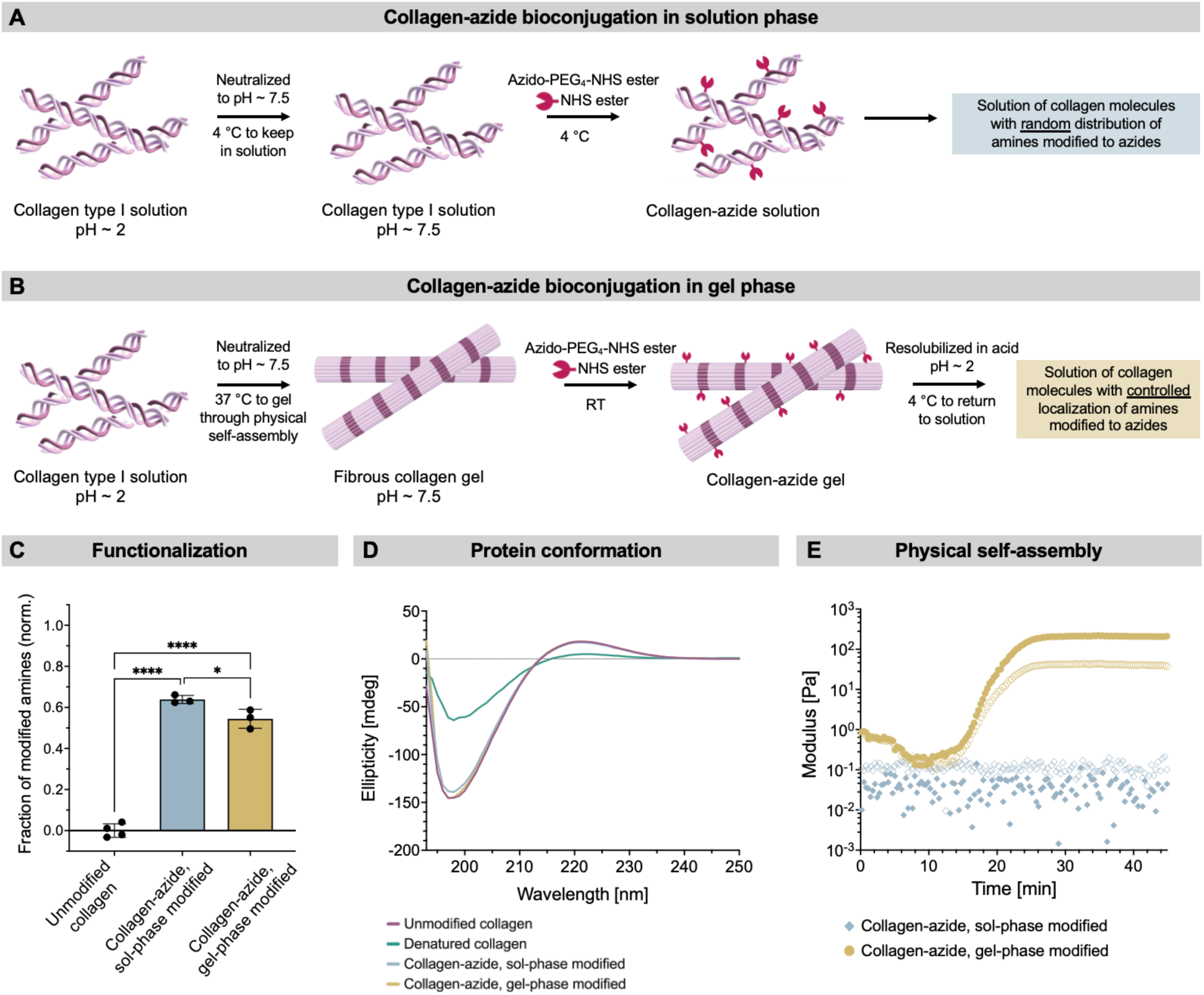
Bioconjugation of collagen with azide functional groups. **(A)** Synthesis approach for collagen-azide bioconjugation in the solution phase. **(B)** Synthesis approach for collagen-azide bioconjugation in the gel phase. **(C)** The collagen-azide degree of functionalization was determined using the relative quantity of free amines before and after the collagen-azide bioconjugation reactions. Normality of the data was confirmed with the Shapiro-Wilk test, and statistical analysis was performed using an ordinary one-way ANOVA with Tukey’s multiple comparisons test. N ≥ 3 independent samples per material condition. Data plotted as mean ± SD. * p = 0.0294, **** p < 0.0001. **(D)** Representative circular dichroism measurements. The collagen-azide solutions retain their triple-helical conformation and exhibit no denaturation. **(E)** Representative gelation curves of 6 mg/mL collagen-azide. Gel-phase modified collagen-azide self-assembles spontaneously from a solution into a hydrogel at a neutral pH upon heating to 37 °C, while sol-phase modified collagen-azide does not spontaneously self-assemble (*i.e.*, remains a solution) under the same conditions. Filled symbols represent the storage modulus (G’), and open symbols represent the loss modulus (G”).

In our first, sol-phase synthesis strategy, the collagen is kept in solution during the bioconjugation reaction, such that all lysine residues on the collagen triple helices are accessible for azide modification (**Figure 1A**). As a result, the chemical modifications in the sol-phase are expected to be non-specifically distributed across the entire collagen molecule. Introducing control over the modification sites, however, could enable greater specificity and tunability of the product’s properties.^31,41^ We therefore adapted a protocol for chemical modification of collagen in the gel-phase,^31^ such that the collagen exists as self-assembled fibrils during the bioconjugation reaction (**Figure 1B**). We hypothesized that this approach would better preserve the fibrous architecture of the collagen, since lysines more involved with collagen self-assembly into fibrils may be protected within the fibrils, while lysines less involved with collagen self-assembly may be surface-exposed and hence preferentially modified.

The modifications of collagen both in the sol-phase and gel-phase were conducted using the same NHS ester reaction chemistry. Since the sol-phase modified collagen-azide remained a neutralized solution throughout the reaction, it was immediately ready for use after purification. In contrast, the gel-phase modified collagen-azide required solubilization for purification and downstream applications, including eventual mixing with cells. Therefore, the collagen-azide gel was reverted to a solution using acid, purified, and then neutralized to restore physiological pH of the solution before use.

Both approaches to synthesize collagen-azide were able to achieve high rates of functionalization, as determined by a fluorescamine assay for free primary amines in collagen before and after the bioconjugation reaction (**Figure 1C**). The degree of amine modification was 64 % for sol-phase modified collagen-azide and 54 % for gel-phase modified collagen-azide. Additionally, both synthesis approaches preserved the secondary structure of native type I collagen, as determined by circular dichroism spectroscopy (**Figure 1D**). The characteristic triple helical conformation of collagen protein was observed for a solution of unmodified collagen (positive control).^42^ Unlike denatured collagen (negative control), solutions of collagen-azide modified either in the sol-phase or gel-phase exhibited similar ellipticity to unmodified collagen, indicating that the introduction of azide groups on the collagen did not significantly alter the secondary protein conformation.

As hypothesized, however, the differences in the physical state of the collagen during the bioconjugation reaction did lead to significant differences in the ability of the collagen-azide solutions to physically self-assemble (**Figure 1E**). The physical self-assembly of collagen into a fibrous hydrogel at a physiological pH (7.4) and temperature (37 °C) is characteristic of native, unmodified PHYS collagen^43,44^ (**Figure S1A**). Solutions of neutralized, 6 mg/mL collagen-azide modified either in the sol-phase or gel-phase were characterized with rheometry for their ability to self-assemble into a hydrogel when heated from 4 °C to 37 °C. No SPAAC crosslinker was added to the solutions, to ensure that any observed gelation was due solely to self-assembly of the collagen-azide. As previously described for sol-phase modified collagen-azide,^14^ the solution did not self-assemble into a hydrogel; the collagen self-assembly process is disrupted by the random distribution of chemical modifications when conducted in the sol-phase.^31^ In contrast, the gel-phase modified collagen-azide retained its ability to self-assemble from a solution into a hydrogel, presumably due to the more controlled localization of azide chemical modifications. Therefore, the collagen-azide synthesis approach in the gel-phase allows the collagen to both be functionalized with azides while preserving the characteristic self-assembling nature of native type I collagen.

We next sought to explore how these two different azide-modified collagens could be blended with unmodified PHYS collagen to form hydrogels crosslinked both through SPAAC covalent crosslinking and physical self-assembly. Depending on whether the collagen-azide for the SPAAC collagen was modified in the sol-phase or gel-phase, distinctly different microstructures were observed (**Figure 2**). In both cases, we formulated the hydrogels to contain 4 mg/mL unmodified PHYS collagen, 2 mg/mL collagen-azide (either modified in sol-phase or gel-phase), and 4 mg/mL PEG-DBCO crosslinker. To compare the microstructures of the resultant hydrogels, we visualized the self-assembled collagen fibrils using their characteristic second harmonic generation (SHG) signal, and the SPAAC-crosslinked collagen using the fluorescence signal from Alexa Fluor 488 (AF488) fluorophores tagged onto pendant azide groups.

**Figure 2.**
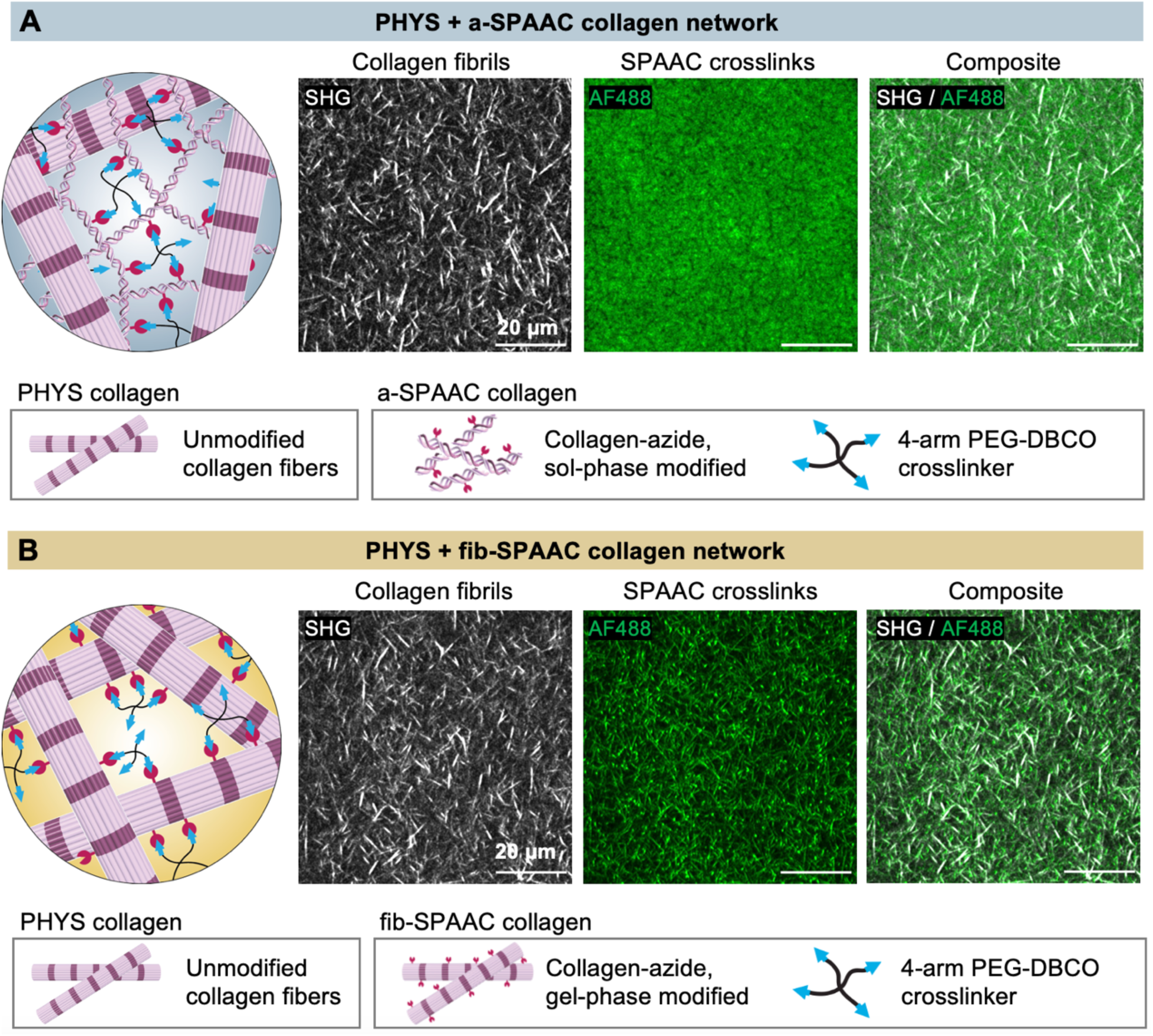
Localization of SPAAC crosslinking sites relative to self-assembled collagen fibrils. The hydrogel formulations are 4 mg/mL unmodified collagen, 2 mg/mL collagen-azide (either sol-phase or gel-phase modified), and 4 mg/mL PEG-DBCO crosslinker. **(A)** PHYS + a-SPAAC collagen (with sol-phase modified collagen-azide) forms an interpenetrating network of fibrous, self-assembled PHYS collagen and amorphous, covalently-crosslinked a-SPAAC collagen. The SHG signals for collagen fibrils and fluorescence signals for SPAAC-crosslinked collagen do not appear to overlap, suggesting that the networks are distinct from each other. **(B)** PHYS + fib-SPAAC collagen (with gel-phase modified collagen-azide) forms a fibrous network in which the sites for SPAAC crosslinking are localized to the fibrils rather than amorphously distributed in the void spaces between them. The SHG signals for collagen fibrils and fluorescence signals for SPAAC-crosslinked collagen appear to overlap, suggesting that the gel-phase modified collagen-azide incorporates into the collagen fibrils with the unmodified collagen.

PHYS collagen on its own self-assembles into a fully fibrillar hydrogel (**Figure S1B**). When sol-phase modified collagen-azide was added to form SPAAC-crosslinked collagen, the SPAAC collagen network was amorphous (a-SPAAC), yielding an interpenetrating network with both fibrillar (from PHYS collagen) and amorphous (from a-SPAAC collagen) constituent networks^14^ (**Figure 2A**). In this case, the signal from the a-SPAAC collagen network appeared to be excluded from the collagen fibrils, consistent with the inability of sol-phase modified collagen-azide to self-assemble into fibrils. Therefore, when mixed with PHYS collagen, the a-SPAAC collagen formed a seemingly independent amorphous network localized to the interfibrillar voids between the PHYS collagen fibrils.

On the other hand, for a hydrogel with the same formulation of components but using collagen-azide modified in gel-phase instead of sol-phase, the SPAAC collagen network was fibrous (fib-SPAAC) rather than amorphous (**Figure 2B**). The signal from the fib-SPAAC collagen appeared to overlap with the SHG signal for total fibrils in the network, indicating that PHYS + fib-SPAAC collagen formed a single fibrous network instead of an interpenetrating network with distinct amorphous and fibrous constituent networks. In this way, by simply changing the physical state of the collagen during the bioconjugation reaction with azides (Figure 1), we demonstrated control over the localization of SPAAC crosslinks within collagen hydrogels (Figure 2). Notably, PHYS + fib-SPAAC collagen offers the possibility to now incorporate sites for SPAAC crosslinking on the native fibrillar architecture of type I collagen.

### 3.2. Sequential fibrillar self-assembly and covalent crosslinking

Fibrous hydrogels with open, interfibrillar voids, such as those observed in the PHYS + fib-SPAAC collagen, have been widely adapted for cell encapsulation. Cell spreading and migration can be enhanced in fibrous hydrogels with voids, and these cell responses are directly impacted by the fiber geometry.^7,8,45^ Thus, we next examined different processing conditions for PHYS + fib-SPAAC collagen to observe their impact on fiber morphology compared to conventional, unmodified collagen hydrogels.

When all hydrogel precursor components for PHYS + fib-SPAAC collagen (*i.e.*, unmodified collagen, gel-phase modified collagen-azide, and PEG-DBCO crosslinker) are mixed together simultaneously, the collagen undergoes concurrent self-assembly into fibrils and covalent SPAAC crosslinking. Concurrent collagen self-assembly and chemical crosslinking is commonly referred to as “simultaneous” processing or gelation.^46^ However, the chemical crosslinking may interfere with the physical self-assembly process by restricting molecular motion, reducing the efficacy of collagen fibril production.^14^ We therefore investigated initiating the fibrillar self-assembly and SPAAC crosslinking either simultaneously or sequentially (**Figure 3**). The sequential formation of networks can often offer greater tunability over the hydrogel microstructure and physical properties.^14,47^

**Figure 3.**
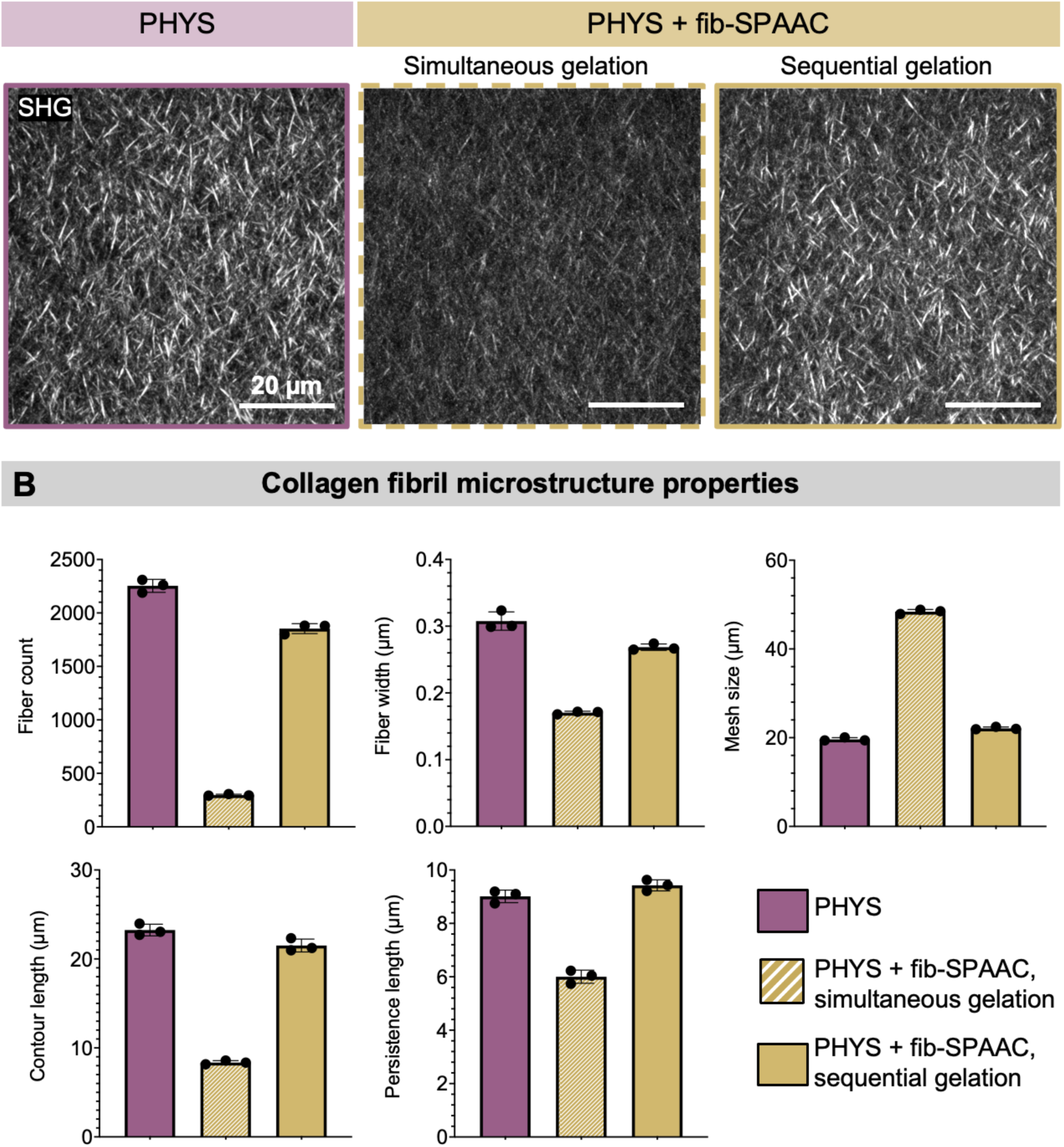
Comparison of fibrillar microstructures for PHYS and PHYS + fib-SPAAC collagen hydrogels. **(A)** Representative SHG images show the fibrillar nature of PHYS collagen, PHYS + fib-SPAAC collagen with simultaneous gelation, and PHYS + fib-SPAAC collagen with sequential gelation. All conditions had a total collagen concentration of 6 mg/mL. For sequential gelation, the PEG-DBCO crosslinker for the SPAAC reaction was introduced only after fibrillar self-assembly was complete. **(B)** The PHYS + fib-SPAAC collagen hydrogels prepared with simultaneous gelation had fewer, thinner, and shorter fibers than PHYS collagen alone. The PHYS + fib-SPAAC collagen hydrogels prepared with sequential gelation had a fibrillar network more closely resembling that of PHYS collagen alone. N = 3 independent samples per material condition. Data plotted as mean ± SD. All groups are statistically different except for comparison between the persistence length of PHYS and PHYS + fib-SPAAC with sequential gelation; statistical analyses presented in Table S1.

To initiate the fibrillar self-assembly and SPAAC crosslinking sequentially, we first allowed a solution of 4 mg/mL unmodified collagen and 2 mg/mL gel-phase modified collagen-azide to undergo physical self-assembly at neutral pH and 37 °C for 1 h. Since no PEG-DBCO crosslinker was present at this stage, no SPAAC crosslinking could proceed. We confirmed with rheometry that the unmodified collagen with gel-phase modified collagen-azide reached a plateau modulus within 30 min, indicating full self-assembly (**Figure S2**). After the collagen self-assembly was complete, we introduced the PEG-DBCO crosslinker by diffusion into the hydrogel, triggering the SPAAC reaction with azides on the collagen for covalent crosslinking.

We visualized the resultant fibrous microstructures using SHG microscopy (**Figure 3A**) and quantitatively analyzed key network parameters—fiber count, width, contour length, persistence length, and network mesh size—using a Fiber Finding Algorithm that traces all fibrils within an imaged volume^14,35^ (**Figure 3B**). We compared the PHYS + fib-SPAAC collagen hydrogels to a PHYS hydrogel with 6 mg/mL unmodified collagen; this maintains the same total collagen concentration, which is a strong influencer of collagen fibril structure.^48^ The PHYS collagen hydrogel therefore served as a positive control for collagen hydrogels formed solely through physical self-assembly, without SPAAC crosslinking.

For PHYS + fib-SPAAC collagen, the simultaneous gelation approach indeed led to competitive interactions between fibrillar self-assembly and SPAAC crosslinking. Compared to PHYS collagen, the PHYS + fib-SPAAC collagen hydrogels prepared with simultaneous gelation had a significant increase in network mesh size and decrease in fiber count, width, contour length, and persistence length (statistical significance values presented in **Table S1**). In contrast, when PHYS + fib-SPAAC collagen was prepared with sequential gelation, the fibrillar network more closely resembled that of PHYS collagen. Therefore, the sequential gelation approach to prepare PHYS + fib-SPAAC collagen preserves the robust fibrillar architecture of conventional PHYS collagen while also incorporating SPAAC crosslinks. All following work on PHYS + fib-SPAAC collagen hydrogels was conducted using the sequential gelation approach.

After examining how the timing of PEG-DBCO crosslinker addition influences the fibrillar microstructure of PHYS + fib-SPAAC hydrogels, we next investigated its impact on the bulk hydrogel’s rheological properties (**Figure 4**). The stiffness and stress relaxation rates of engineered matrices are known to influence the phenotype of encapsulated cells.^49,50^ Specifically, we compared hydrogels before the addition of the PEG-DBCO SPAAC crosslinker (*i.e.*, 4 mg/mL unmodified collagen, 2 mg/mL gel-phase modified collagen-azide) and hydrogels after SPAAC crosslinking (*i.e.*, 4 mg/mL unmodified collagen, 2 mg/mL gel-phase modified collagen-azide, 4 mg/mL PEG-DBCO; prepared with the sequential gelation approach). Interestingly, the stiffness (**Figure 4A**) and stress relaxation behavior (**Figure 4B**) of the hydrogels with and without the SPAAC crosslinker were similar. The storage modulus (G’, energy stored elastically) was ∼250 Pa and the stress relaxation half-time (τ_1/2_, time for the initial peak stress to relax by half) was ∼500 s. The rapid stress-relaxing behavior is characteristic of fibrillar, self-assembled collagen hydrogels and enables key cellular processes such as matrix remodeling, spreading, and migration.^51–53^

**Figure 4.**
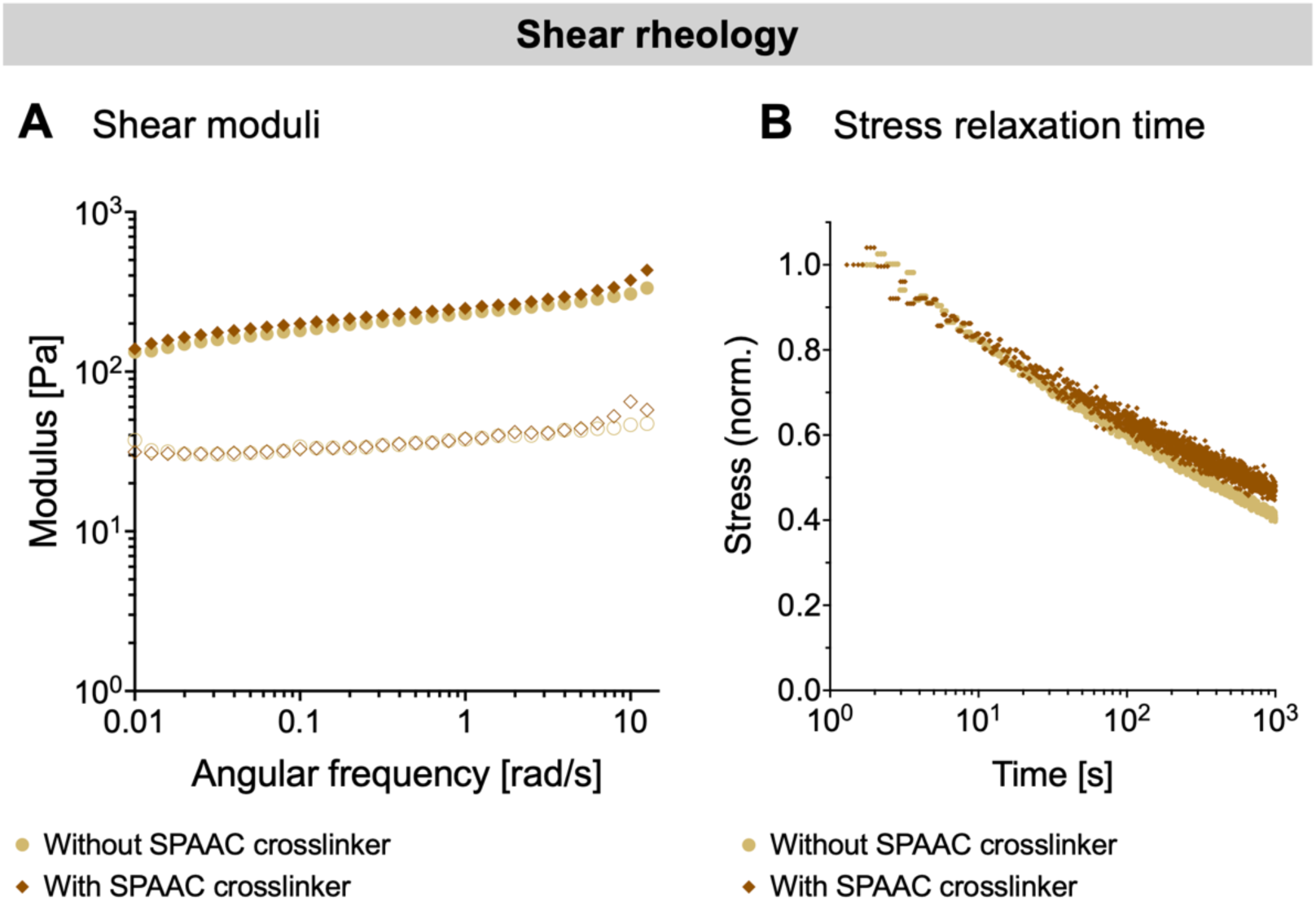
Rheological properties of PHYS + fib-SPAAC collagen hydrogels without and with addition of the SPAAC crosslinker. Conditions without the SPAAC crosslinker were formulated as 4 mg/mL unmodified collagen and 2 mg/mL gel-phase modified collagen. Conditions with the SPAAC crosslinker included 4 mg/mL PEG-DBCO, introduced with the sequential gelation approach. **(A)** Representative hydrogel shear moduli curves. Filled symbols represent the storage modulus (G’), and open symbols represent the loss modulus (G”). **(B)** Representative hydrogel stress relaxation curves.

The similarity of the shear moduli and stress relaxation times between conditions with and without the SPAAC crosslinker suggest that the SPAAC crosslinking is primarily intrafibrillar (*i.e.,* occurring across collagen molecules within a single fibril) rather than interfibrillar (*i.e.,* occurring between adjacent fibrils). Given the molecular weight of the crosslinker (4 arms, 10 kDa total), the maximum distance connecting two crosslinking sites is < 40 nm. This distance is smaller than the observed interfibrillar voids (Figure 3), likely reducing instances in which the crosslinker bridges across adjacent fibrils.

### 3.3. Hydrogel cytocompatibility and stability

Due to the prominence of collagen in the human body, collagen-based hydrogels have broad applications for fabrication of cell-laden engineered tissues including skin, muscle, and connective tissues.^54,55^ In the corneal stroma, for example, collagen accounts for more than 70 % of the dry weight.^56^ The collagen matrix is crucial for the biomechanical properties of the cornea that impart its form and function.^57,58^ Given the global shortage of transplantable donor corneas for patients with corneal blindness,^59^ there is great interest in developing bioengineered corneal tissue as an alternative.^60^ Corneal mesenchymal stromal cells (MSCs) play a key role in corneal regeneration and can be easily isolated and expanded, thus making them promising candidates for incorporation into engineered corneal tissues.^15,32,61,62^ As a demonstrative example of the suitability of PHYS + fib-SPAAC collagen hydrogels to support cell encapsulation, we encapsulated human corneal MSCs within the hydrogels at a density of 0.5 million cells/mL (**Figure 5**). Conventional PHYS collagen hydrogels with the same density of encapsulated corneal MSCs and the same collagen concentration (6 mg/mL) served as the control.

**Figure 5.**
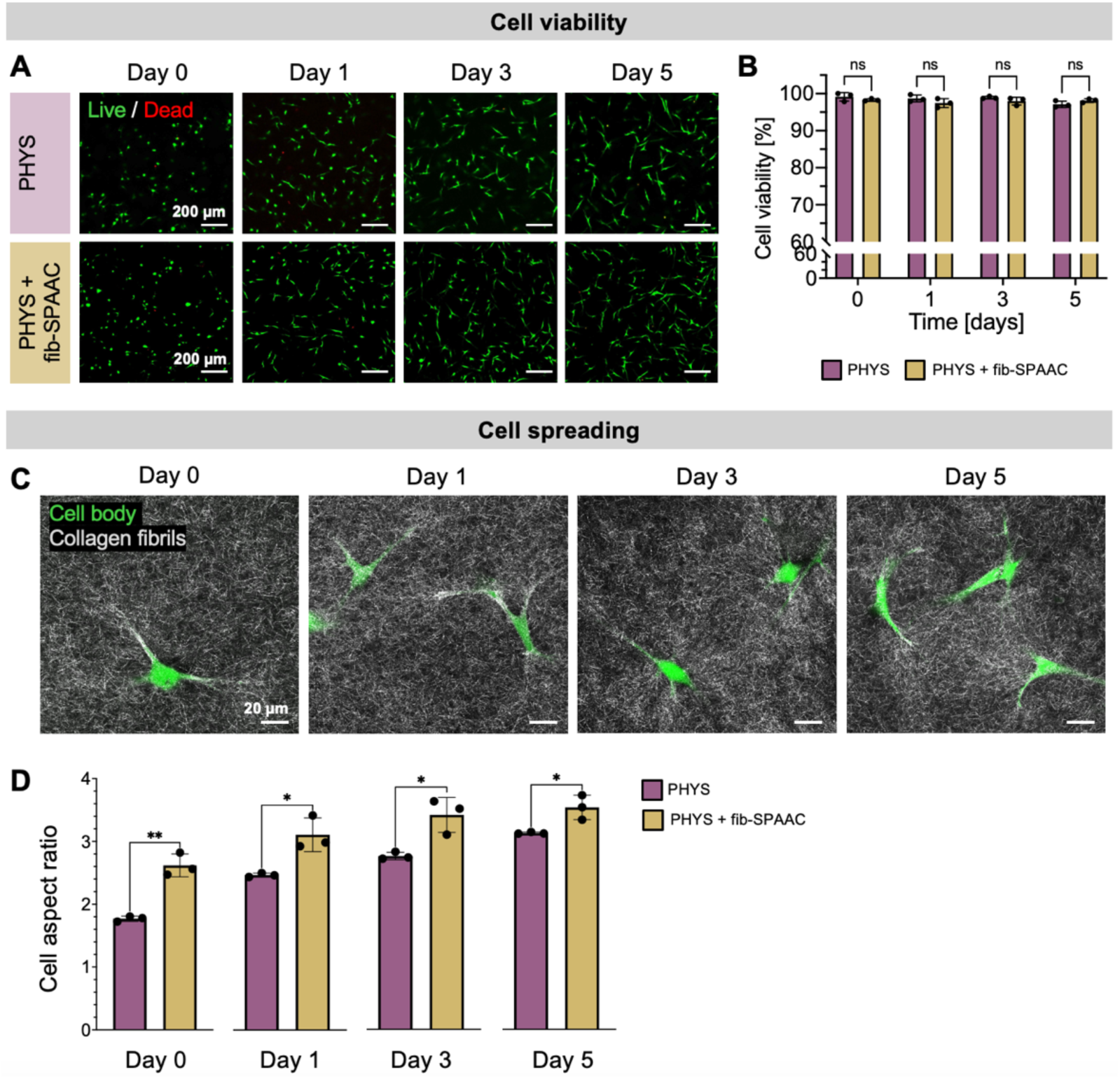
Viability and spreading of encapsulated human corneal MSCs. Human corneal MSCs were encapsulated within PHYS and PHYS + fib-SPAAC collagen hydrogels on Day 0. All material conditions had a total collagen concentration of 6 mg/mL collagen. **(A)** Representative images from Live/Dead cytotoxicity assays of corneal MSCs on Days 0, 1, 3, and 5. **(B)** Corneal MSCs remain highly viable within both PHYS and PHYS + fib-SPAAC collagen over 5 days in culture. **(C)** Representative fluorescence and confocal reflectance images of corneal MSCs within PHYS + fib-SPAAC collagen hydrogels show cell spreading and local interactions of cells with the collagen matrix. **(D)** Quantification of the aspect ratios of cells throughout the culture period. Statistical analyses were performed with unpaired t-tests for each timepoint. In all cases, N = 3 independent gels per material condition and timepoint. Data plotted as mean ± SD. ns = not significant, * p < 0.05, ** p < 0.01.

The corneal MSCs exhibited high viability within PHYS + fib-SPAAC collagen hydrogels over 5 days in culture (**Figure 5A**). For all timepoints, more than 95 % of the cells were alive, similar to the conventional PHYS collagen hydrogel control (**Figure 5B**). These results confirm the high cytocompatibility of the PHYS + fib-SPAAC collagen material and the sequential crosslinking process, which consists of both the collagen physical self-assembly and the covalent SPAAC crosslinking. As the SPAAC chemistry is bioorthogonal, it does not cross-react with chemical functional groups present on the surfaces of cells or in culture medium.^27^ This reaction can therefore proceed in the presence of the living cells that are pre-mixed with the collagen solution prior to gelation.

Cell morphology is also an important indicator of cellular functions.^63,64^ In the native cornea, corneal MSCs adopt spread and elongated morphologies throughout the collagen-rich stromal tissue.^65,66^ We observed that corneal MSCs appeared to spread their cell bodies and remodel the PHYS + fib-SPAAC collagen fibrils in the pericellular regions near cell projections (**Figure 5C**). The extent of cell spreading within the hydrogels was quantified as the average cell aspect ratio (*i.e.*, ratio of the major to minor axis lengths for the best-fit ellipse of each cell) (**Figure 5D**). Previous studies have observed that corneal MSCs in amorphous, SPAAC-crosslinked collagen (a-SPAAC) remain rounded over time.^13–15^ Average aspect ratios of corneal MSCs in a-SPAAC collagen were < 2 on Day 5 in culture.^14^ In contrast, corneal MSCs in PHYS + fib-SPAAC collagen elongated to reach an aspect ratio of ∼ 3.5 on Day 5 in culture. The cells in PHYS + fib-SPAAC collagen also had higher aspect ratios than those in PHYS collagen hydrogels across all timepoints, indicating efficient cell spreading.

Cell-laden engineered tissues should maintain appropriate structural stability for their intended application. Rapid deformation or degradation of an engineered tissue, for example, can compromise its functionality for use as an *in vitro* model or for *in vivo* transplant. Therefore, we next characterized the stability of the PHYS + fib-SPAAC collagen hydrogels compared to conventional PHYS collagen hydrogels to evaluate if the SPAAC crosslinks help to maintain mechanical integrity against cell-induced contraction and enzymatic digestion (**Figure 6**).

**Figure 6.**
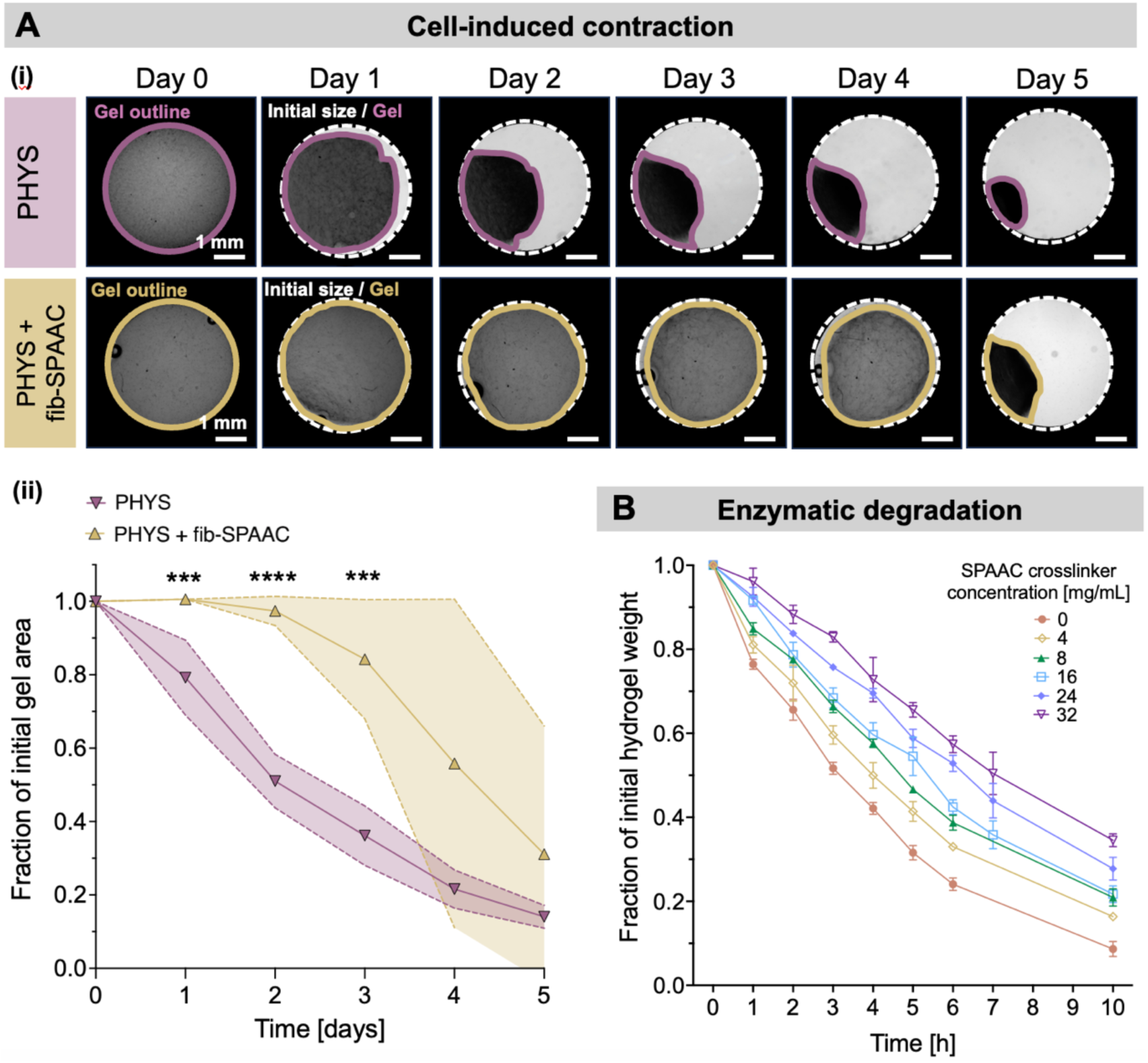
Stability of PHYS + fib-SPAAC collagen hydrogels. All material conditions had a total collagen concentration of 6 mg/mL collagen. **(A)** Stability of hydrogels against cell-induced contraction. All hydrogels were prepared with an initial density of 0.5 million corneal MSCs/mL. (i) Representative brightfield images of the bulk size of PHYS and PHYS + fib-SPAAC collagen hydrogels with encapsulated corneal MSCs over the culture time. Dashed white outlines indicate the edges of the mold, which is the initial size of the hydrogels. Solid colored outlines indicate the edges of the hydrogel at each timepoint. (ii) Quantified rate of collagen hydrogel contraction. Statistical analyses were performed with unpaired t-tests to compare the material conditions at each timepoint. N ≥ 6 independent samples per material condition. Shaded regions represent the standard deviation from the mean. *** p < 0.001, **** p < 0.0001. **(B)** Stability of PHYS + fib-SPAAC collagen hydrogels against 0.5 wt% collagenase. All formulations contained 4 mg/mL unmodified collagen and 2 mg/mL gel-phase modified collagen-azide, and the concentration of PEG-DBCO as the SPAAC crosslinker was varied between 0-32 mg/mL. N = 3 independent samples per material condition. Data plotted as mean ± SD. Statistical analysis for t = 10 h is shown in Table S2.

Like other types of MSCs, corneal MSCs are contractile cells that exert tensions and forces on their surrounding matrix.^13–15,67^ One limitation of conventional PHYS collagen hydrogels is their rapid and severe contraction in size in response to contractile forces generated by encapsulated cells.^10–12^ Corneal MSCs were used here as a demonstrative example of a highly contractile cell type, since hydrogel contraction would not be expected for cell types with low traction force generation.^68^ We compared the effects of corneal MSC-induced contraction between PHYS and PHYS + fib-SPAAC collagen hydrogels over 5 days in culture (**Figure 6A**). All hydrogels were prepared in circular 4-mm diameter molds with the same total collagen concentration (6 mg/mL) and the same cell density (0.5 million corneal MSCs/mL). Brightfield images were taken daily to track the sizes of the hydrogels. While both the cellular PHYS and PHYS + fib-SPAAC collagen hydrogels exhibited some extent of contraction by Day 5, the contraction of PHYS + fib-SPAAC collagen was slower than the PHYS collagen hydrogel, indicating greater structural stability.

Collagen hydrogels are also susceptible to enzymatic degradation mediated primarily by collagenases, a subgroup of metalloproteinases that cleave the collagen’s peptide backbone.^69^ In native healthy tissues, collagenase activity is regulated to enable controlled ECM remodeling during development, wound healing, and homeostasis.^69^ Excessive degradation of collagen-based engineered tissues, however, can lead to premature structural failure.^70^ It is therefore useful to tailor the degradation rate of tissue-engineered constructs to match the specific application, in order to achieve a balance between ECM remodeling and sustained structural integrity.^17,70^ We performed an accelerated enzymatic digestion assay on PHYS + fib-SPAAC collagen hydrogels by exposing them to 0.5 wt% collagenase. The PHYS + fib-SPAAC collagen hydrogels exhibit tunable degradation behavior when the concentration of the PEG-DBCO molecule for SPAAC crosslinking is adjusted (**Figure 6B**). Even for the lowest concentration of PEG-DBCO tested (4 mg/mL, identical to the formulation used for all other experiments), the rate of enzymatic degradation was significantly decreased (statistical significance shown in **Table S2**). With increasing amounts of PEG-DBCO, the rate of enzymatic degradation is further slowed. Therefore, PHYS + fib-SPAAC collagen is a tunable and versatile hydrogel platform, combining the biostructural properties of fibrillar networks with enhanced stability provided by SPAAC crosslinks localized along the fibrils.

## 4. Conclusions

In summary, we developed a collagen hydrogel that not only self-assembles into a fibrillar network but also presents azide groups along the collagen fibrils for SPAAC crosslinking. By modifying collagen with azides in the gel-phase rather than sol-phase, we preserved its ability to self-assemble into fibrils, unlike previous approaches.^13–15,29,30^ The PHYS + fib-SPAAC collagen hydrogel maintains similar fibrillar network properties as PHYS collagen alone when prepared with a sequential gelation technique, in which the modified collagen is allowed to fully self-assemble into fibrils before addition of the SPAAC crosslinker. The PHYS + fib-SPAAC collagen hydrogels support the high viability and spreading of encapsulated corneal MSCs, demonstrating the cytocompatibility of the material system and crosslinking procedure. Finally, PHYS + fib-SPAAC collagen hydrogels exhibit enhanced stability, with greater resistance to cell-induced contraction than conventional PHYS collagen hydrogels and a tunable enzymatic degradation rate by modulating the SPAAC crosslinker concentration.

Future work could vary the PHYS + fib-SPAAC collagen concentrations and ratio of PHYS to fib-SPAAC collagen. Using a PEG-DBCO crosslinker with longer PEG arms may also increase the distance over which crosslinking occurs, which could increase the propensity for interfibrillar connections and be also used to tune stiffness, stress relaxation times, and bulk hydrogel stability. Bespoke formulations with suitable biomechanical and structural properties could therefore be selected for specific target tissues and applications. Furthermore, the PHYS + fib-SPAAC collagen could be applied as a hydrogel bioink for embedded 3D bioprinting,^71,72^ in which the SPAAC crosslinker diffuses from a support bath into the bioink for covalent crosslinking.^13,73^ Finally, while we demonstrated use of the azides functionalized onto collagen for crosslinking, SPAAC chemistry could instead be used to tether bioactive molecules such as growth factors or peptide ligands onto the collagen fibrils.^74,75^ For example, pendant azides on hydrogels implanted *in vivo* can be used as targetable and replenishable drug depots, through the circulation of drugs conjugated with strained alkynes.^76,77^ Overall, the PHYS + fib-SPAAC collagen hydrogel presents a versatile and biofunctional platform that preserves the native fibrillar architecture of collagen while enabling bioorthogonal, covalent chemistry, offering broad potential for tissue engineering, biofabrication, and drug delivery applications.

## Supporting Information

Additional experimental details, including rheological characterizations, hydrogel microstructure images, and statistical significance comparisons between fibrillar network properties and degradation rates.

## Acknowledgements

The authors thank Prof. Possu Huang for CD spectrometer access, Prof. Ninna Rossen for helpful discussions about the Fiber Finding Analysis, Dr. Chris Lindsay and Dr. Sarah Hull for helpful discussions about the collagen-azide bioconjugation protocols, and Dr. Philipp Fisch for assistance with schematics. The authors acknowledge funding support from the National Science Foundation including DGE-165618 (L.G.B., C.M.L.), DMR-2103812 (S.C.H.), DMR-2427971 (S.C.H.), and CBET-2033302 (S.C.H); the National Institutes of Health including F31-EY034785 (L.G.B), P30-EY026877 (D.M.), R01-EY033363 (D.M.), and R01-EY035697 (D.M., S.C.H.); the ARCS Foundation Scholarship (L.G.B.); the Stanford Bio-X Interdisciplinary Graduate Fellowship (C.M.L., B.C.); the Swiss National Science Foundation including P500PN210723 (F.C.); the Stanford Knight-Hennessy Scholars Program (B.C.); the American Heart Association including 24PRE1191604 (N.d.P.N.); the Gerald J. Lieberman Fellowship (N.d.P.N.); the Stanford Graduate Fellowship in Science and Engineering (D.S.); a departmental core grant from Research to Prevent Blindness (D.M.); and the Advanced Research Projects Agency for Health including ARPA-H HEART (S.C.H.).

## Data Availability

All data needed to evaluate the conclusions in the paper are present in the paper and/or the Supplementary Information.

## Table of Contents Graphic

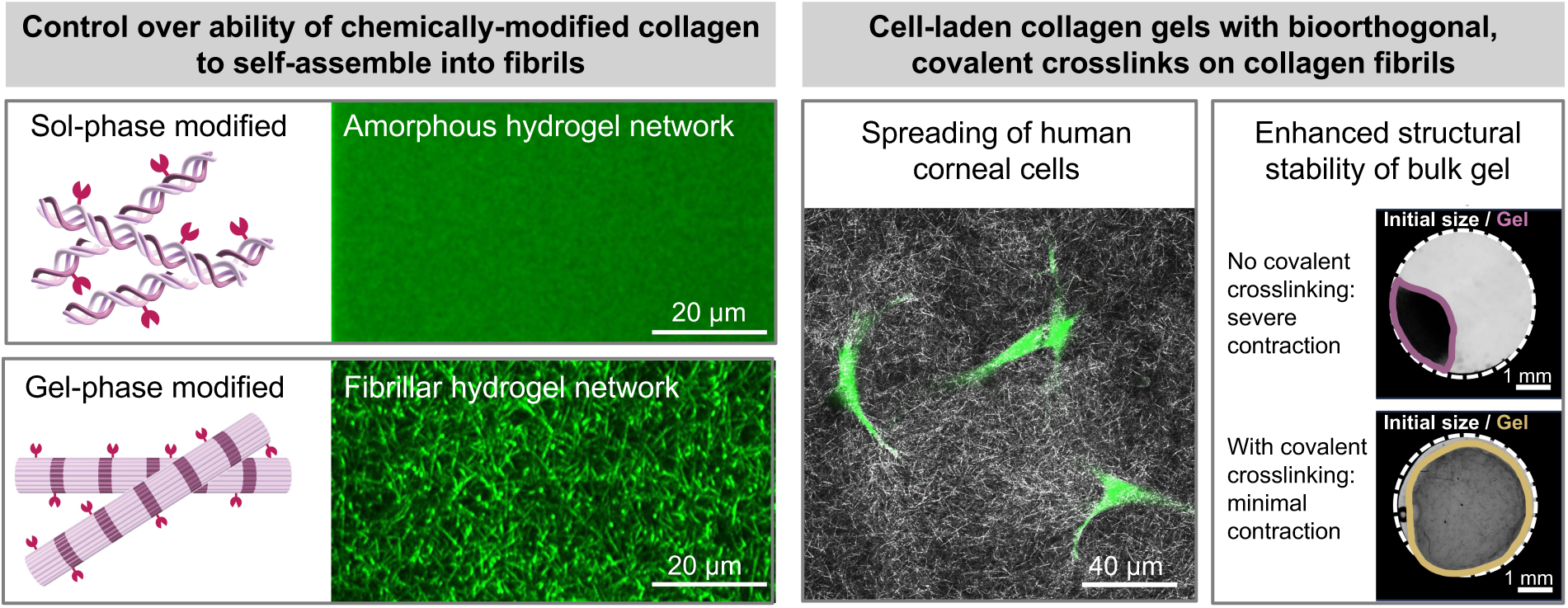

