## Supplementary Information for "Reinforcement of Fibrillar Collagen Hydrogels with Bioorthogonal Covalent Crosslinks"

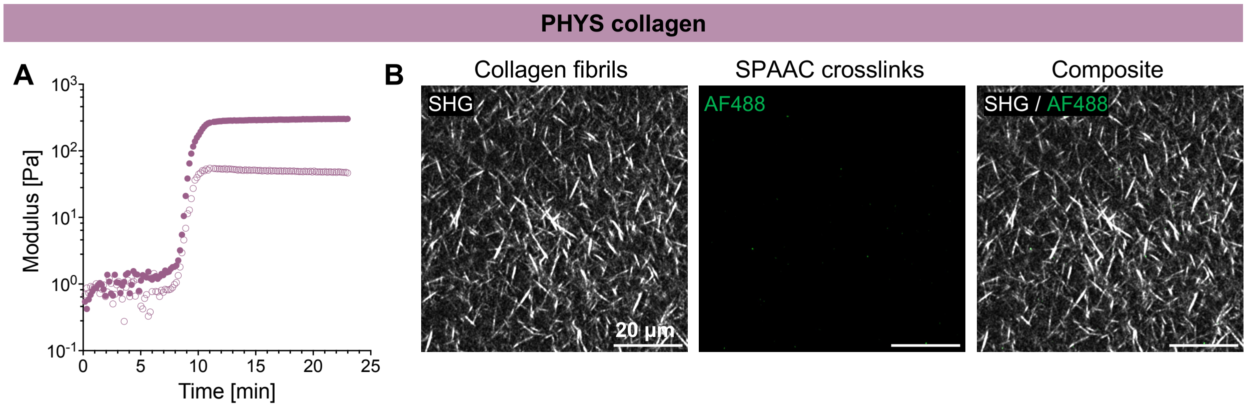


**Figure S1.** Gelation kinetics and structural properties of 6 mg/mL PHYS collagen hydrogels. **(A)** PHYS collagen at a neutral pH spontaneously self-assembles into a hydrogel within 10 min upon heating to 37 °C. Filled symbols represent the storage modulus (G'), and open symbols represent the loss modulus (G"). **(B)** PHYS collagen hydrogels consist of a fibrillar network with no covalent SPAAC-crosslinked network. This serves as a negative control for fluorescently labelling sites on collagen available for SPAAC crosslinking.

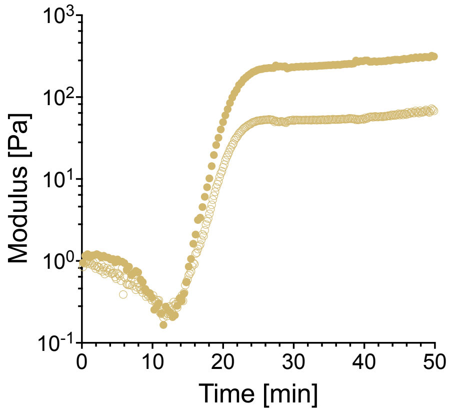


**Figure S2.** Gelation kinetics of 4 mg/mL unmodified collagen with 2 mg/mL gel-phase modified collagen-azide. The collagen formulation spontaneously self-assembles into a hydrogel at a neutral pH within 30 min upon heating to 37°C. Filled symbols represent the storage modulus (G'), and open symbols represent the loss modulus (G").

**Table S1.** Statistical comparison of fibril network properties between PHYS collagen and PHYS + fib-SPAAC collagen with either simultaneous or sequential gelation of the PHYS and fib-SPAAC networks. N = 3 independent samples per material condition. Statistical analyses were performed using an ordinary one-way ANOVA with Tukey's multiple comparisons test.

| | **PHYS** *vs.* **PHYS + fib-SPAAC, sim.** | **PHYS** *vs.* **PHYS + fib-SPAAC, seq.** | **PHYS + fib-SPAAC, sim.** *vs.* **PHYS + fib-SPAAC, seq.** |
|---|---|---|---|
| **Contour length** | <0.0001 (****) | 0.0240 (*) | <0.0001 (****) |
| **Persistence length** | <0.0001 (****) | 0.1501 (ns) | <0.0001 (****) |
| **Mesh size** | <0.0001 (****) | 0.0007 (***) | <0.0001 (****) |
| **Fiber count** | <0.0001 (****) | <0.0001 (****) | <0.0001 (****) |
| **Fiber width** | <0.0001 (****) | 0.0032 (**) | <0.0001 (****) |

**Table S2.** Statistical comparison of the fraction of hydrogel remaining after 10 h of 0.5 wt% collagenase treatment, for varying concentrations of PEG-DBCO crosslinker. N = 3 independent samples per material condition. Statistical analyses were performed using an ordinary one-way ANOVA with Tukey's multiple comparisons test.

| **Concentration of PEG-DBCO [mg/mL]** | **0** | **4** | **8** | **16** | **24** | **32** |
|---|---|---|---|---|---|---|
| **0** | - | 0.0033 (**) | <0.0001 (****) | <0.0001 (****) | <0.0001 (****) | 0.0001 (****) |
| **4** | - | - | 0.0983 (ns) | 0.0433 (*) | <0.0001 (****) | <0.0001 (****) |
| **8** | - | - | - | 0.9954 (ns) | 0.0085 (**) | <0.0001 (****) |
| **16** | - | - | - | - | 0.0195 (*) | <0.0001 (****) |
| **24** | - | - | - | - | - | 0.0091 (**) |
| **32** | - | - | - | - | - | - |
